# Characterization of *Neisseria gonorrhoeae* colonization of macrophages under distinct polarization states and nutrients environment

**DOI:** 10.1101/2024.02.08.579566

**Authors:** María Dolores Juárez Rodríguez, Madison Marquette, Reneau Youngblood, Nilu Dhungel, Ascención Torres Escobar, Stanimir Ivanov, Ana-Maria Dragoi

## Abstract

*Neisseria gonorrhoeae (Ng)* is a uniquely adapted human pathogen and the etiological agent of gonorrhea, a sexually transmitted disease. *Ng* has developed numerous mechanisms to avoid and actively suppress innate and adaptive immune responses. *Ng* successfully colonizes and establishes topologically distinct colonies in human macrophages and avoids phagocytic killing. During colonization, *Ng* manipulates the actin cytoskeleton to invade and create an intracellular niche supportive of bacterial replication. The cellular reservoir(s) supporting bacterial replication and persistence in gonorrhea infections are poorly defined. The manner in which gonococci colonize macrophages points to this innate immune phagocyte as a strong candidate for a cellular niche during natural infection. Here we investigate whether nutrients availability and immunological polarization alter macrophage colonization by *Ng*. Differentiation of macrophages in pro-inflammatory (M1-like) and tolerogenic (M2-like) phenotypes prior to infection reveals that *Ng* can invade macrophages in all activation states, albeit with lower efficiency in M1-like macrophages. These results suggest that during natural infection, bacteria could invade and grow within macrophages regardless of the nutrients availability and the macrophage immune activation status.

## Introduction

Gonorrhea, a sexually transmitted infection (STI) caused by the obligate human pathogen *Neisseria gonorrhoeae* (*Ng*) is rising worldwide with > 80 million new cases annually (1, 2). *Ng* mainly colonizes and infects the urogenital tract (3). In men, gonorrhea mostly manifests as symptomatic urethritis, while in women the infection can remain asymptomatic making detection and early treatment difficult. A significant number of women (15%) develop ascending pelvic inflammatory disease with serious complications including chronic pelvic pain, infertility, increased risk of ectopic pregnancy, and endometriosis (4, 5, 6). One clinical problem is the widespread rapid emergence of multi-drug resistant strains, which limits therapeutic options. In 2019, CDC classified drug-resistant *Ng* as an urgent threat. Another significant challenge is vaccine development, due to the diverse immune evasion mechanisms utilized by *Ng* to avoid killing and suppress immunological responses (3, 7).

The localized innate immune response at the sites of bacterial colonization drives the immunopathology central to gonorrhea disease (3, 8, 9). Macrophages and neutrophils are key players in mucosal immune defenses (10). In symptomatic gonorrhea, monitoring exudate secretions for neutrophils harboring gonococci is a classical early diagnostic test (3). Thus, historically, neutrophils have been the research focus (11, 12, 13, 14, 15, 16) and many aspects of *Ng* interaction with neutrophils have been elucidated, including the observation that neutrophils can support limited bacterial replication *ex vivo* (16). New data, however, demonstrated that macrophages have a high capacity to support *Ng* replication (>2−3 orders of magnitude in 8hrs) in nutrient-depleted environments for >24 hrs (17, 18). Exudate secretions in acute gonorrhea besides neutrophils and exfoliated epithelial cells also contain macrophages (9). Furthermore, macrophages are present throughout the genitourinary mucosa, representing about 10% of all mucosa-resident leukocytes (8, 19), and are recruited to and infiltrate the uterine mucosa in murine models of vaginal gonorrhea (20, 21). Thus, macrophages could be one cellular niche for *Ng* replication and persistence in human infections. Indeed, we showed that *Ng* can colonize and invade macrophages forming both surface-associated as well as intracellular colonies (18). The invasion process is carried out by a novel mechanism via FMNL3-regulated actin-dependent membrane morphogenesis, which was observed for both human differentiated U937 macrophages as well as primary human monocyte-derived macrophages (hMDMs). Additionally, gonococci can inhibit apoptosis in human macrophages, and have been shown to elicit secretion of both inflammatory and tolerogenic cytokines by macrophages (17, 22, 23). The emerging data argue for further investigation of macrophages as a cellular niche and regulators of immunological responses in gonorrhea.

*Ng* colonizes macrophages in a manner that facilitates immune evasion. We have shown that an early step in invasion is the establishment of a surface microcolony, which entails attachment, inhibition of canonical phagocytosis, and bacterial replication on the surface of the host cell (18). Some colonies remain surface-associated, while others trigger membrane remodeling and formation of an invasion platform where the actin nucleating factor FMNL3 is recruited and promotes actin polymerization and colony uptake (18). During invasion, the colony stretches from the surface of the host cell into a plasma membrane-derived membrane-bound compartment that cradles a fraction of the colony. The simultaneous existence of surface-associate, intracellular, and partially internalized hybrid bacterial colonies would benefit immune evasion. This adaptation to multiple topologically distinct niches is a complex evolutionary solution counteracting multiple distinct host defense responses. A surface-associated niche protects *Ng* from several cell-autonomous defense responses against intracellular pathogens (such as lysosomal degradation, autophagy, and host cell death) (24, 25, 26), whereas an intracellular niche safeguards against humoral defenses targeting extracellular bacteria (such as complement, circulating antibodies and anti-microbial peptides) (27). Indeed, when macrophages invasion was blocked by cytochalasin D or in FMNL3-depleted cells, the number of bacteria protected from antibiotic killing decreased by ∼70% as determined by CFU assays (18). Thus, the FMNL3-mediated gonococcal internalization by human macrophages protects gonococci from extracellular microbiocidal activities.

The impact of nutrients availability and macrophage immune polarization on gonococci capacity to colonize, invade and replicate within human macrophages is investigated here. The extracellular micronutrients in the urogenital mucosa vary based on gender (28, 29, 30, 31) and over time during the reproductive cycle in women (32). Thus, it is important to establish whether nutrients limitation impacts macrophage colonization and invasion by gonococci. In addition, macrophages can undergo phenotypic polarization in response to the environmental cues to carry out tissue surveillance and homeostasis functions (33, 34). The distinct polarization states can impact macrophage-driven immunological responses to infection (35, 36, 37). The classically activated M1 macrophages have enhanced microbiocidal capacity, secrete pro-inflammatory/T helper 1 (Th1)-promoting cytokines (IL-12, IL-1β, IL-6, and TNF-α), and mediate host defense against bacterial, viral and protozoal pathogens (36). The alternatively activated M2 macrophages mediate tissue repair and exhibit a more tolerogenic immunological profile by preferential secretion of anti-inflammatory mediators (IL-10 and TGF-β) (38). Polarization towards the M1 state can be elicited by stimulation with IFN-γ and lipopolysaccharide (LPS), while M2 polarization is triggered by stimulation with IL-4, IL-10 and IL-13 (39). Because gonococci are likely to encounter macrophages in distinct polarization states in symptomatic *vs.* asymptomatic infections, it is important to determine the impact on colonization and invasion.

## Materials and Methods

### Bacterial strains and culture conditions

*Neisseria gonorrhoea*e FA1090, was a gift from Dr. Hank Seifert (Northwestern University). FA1090-LuxR strain was generated as detailed below. Bacteria were cultured on gonococcal medium base (GCB) agar (Criterion) with Kellogg’s supplements (40) at 37°C and 5% CO_2_ for approximately 16 hrs. For infections, 4 to 6 piliated Opa+ colonies as determined by colony morphology were pooled, streaked on GCB agar (1×1cm patch) and grown for another 16 hrs. Patches were collected and resuspended in PBSG (PBS supplemented with 7.5 mM Glucose, 0.9 mM CaCl_2_, and 0.7 mM MgCl_2_). The number of bacteria was determined by OD_600_ measurements. Inocula for infections were prepared in PBSG, RPMI or RPMI containing inactivated human serum (10%) and the number of bacteria was confirmed by plating dilutions from the inoculum on GCB agar. *Legionella pneumophila* serogroup 1 strain JR32 Δ*flaA luxR* was used in this study (41). The *Legionella* strain was grown on charcoal yeast extract (CYE) plates composed of 1% yeast extract, 1%N-(2-acetamido)-2-aminoethanesulphonic acid (ACES; pH 6.9), 3.3 mM l-cysteine, 0.33 mM Fe(NO3)_3_, 1.5% agar, 0.2% activated charcoal] (42). For macrophage infections *Legionella* was obtained from day 2 heavy patches grown on CYE plates and was resuspended to OD_600_ of 0.5 U in 1 ml AYE, placed in 15-ml glass culture tubes, and cultivated aerobically 24 to 26 hrs with continuous shaking (175 rpm) at 37°C until early stationary phase was reached (OD_600_ range of 2.0 to 3.0 U). For propagation and plating of *Escherichia coli*, Luria-Bertani (LB) broth and LB agar (LB broth with 1.5% agar) were used. As needed, the culture media were supplemented with 2 µg/ml erythromycin.

### Construction of the plasmid pMR32-MCS*-luxCDABE*

The construction of the plasmid pMR32-MCS-*luxCDABE* for the generation of *Neisseria gonorrhoeae* FA1090-LuxR luminescent strain was performed in two steps: (1) A 95-bp DNA fragment containing a multi-cloning site flanked by cohesive ends for PmeI and ApaI restriction enzymes, was generated by the annealing of the following primers:

pMR32MCS_F (5′AAACTCACTAGTCATGTCGACTTCGAATTCGTCCATTCCTGCAGGTCTGAGCTCTATCCTAGGATGCCGTCCGAACCTTCAGACGGCATTGGGCC 3′)

pMR32MCS_R (5′CAATGCCGTCTGAAGGTTCGGACGGCATCCTAGGATAGAGCTCAGACCTGCAGGAATGGACGAATTCGAAGTCGACATGACTAGTGAGTTT 3′).

The resulting DNA product was phosphorylated and cloned into the pMR32 plasmid (43) digested with PmeI and ApaI to obtain the pMR32-MCS plasmid. (2) The *Photorharddus luminescens luxCDABE* operon was PCR amplified using as a template the pXen-13 plasmid (Xenogen Bioware) with the following primer set:

lux_F (5′ TATGGGGAAGTCGACTTGGAGGATACGTATGACTAAAAAAATTTC 3′)

lux_R (5′ GAATTAACGAGCTCGAATACCTGCAGGTCATCAACTATCAAAC 3′)

The 5857-bp amplicon was digested with SalI and SacI and cloned into the pMR32-MCS plasmid digested with the same enzymes to generate the pMR32-MCS-*luxCDABE* plasmid that harbors the *Photorharddus luminescens luxCDABE* operon under the control of *Neisseria gonorrhoeae opaB* promoter.

### Generation of *Neisseria gonorrhoeae* FA1090-LuxR luminescent strain

*Neisseria gonorrhoeae* FA1090 was grown from frozen stocks onto the GCB agar (Criterion) with 1% Kellogg’s supplements at 37°C and 5% CO_2_ for 16 hrs. From bacteria grown on GCB agar, 5 piliated colonies were collected and pooled in GC Broth (1.5% Difco proteose peptone no. 3, 0.4% K_2_HPO_4_, 0.1% KH_2_PO_4_, 0.1% NaCl, pH of 7.2, with 1% Kellogg’s supplements and 0.042% NaHCO_3_), and were naturally transformed with 1 µg of pMR32-MCS-*luxCDABE* plasmid DNA onto GCB agar. The plate was incubated at 37°C with 5% CO_2_, for 16 h. The next day, the bacteria were harvested from the plate, resuspended in GC Broth, seeded onto GCB agar plates containing 2µg/ml erythromycin, and incubated at 37°C with 5% CO_2_, for 16h. The erythromycin-resistant colonies were collected and analyzed by PCR amplification with the following primer set:

F_ermC (5′-GAACATGATAATATCTTTGAAATCGGCTCAG -3′)

R_ermC (5′ -CTTGTATTCTTTGTTAACCCATTTCATAACG-3′).

Allelic exchange was confirmed in individual colonies by kanamycin-sensitivity, erythromycin-resistance, PCR amplification of the 525-bp fragment and bioluminescence.

### Ratio of CFUs to bioluminescence

*Neisseria gonorrhoeae* FA1090-LuxR strain was grown on GCB agar plates supplemented with Erythromycin at 2 µg/ml, harvested in GC Broth, and resuspended at an OD_600_ = 0.5 U in GC broth. Then, 2-fold serial dilutions were made in the same medium and 200 µl aliquots of each dilution were distributed in triplicate in a white-wall clear bottom 96-well assay plate (Corning). The OD_600_ and luminescence data were acquired in a luminescence plate reader (BioTek Synergy H1, Agilent). The number of bacterial colony-forming units (CFUs) was determined by plating serial dilutions from which bioluminescence light output per OD_600_ unit or per CFU were calculated.

### Axenic growth kinetics of *Neisseria gonorrhoeae* strains

Bacteria grown on GCB agar were harvested and resuspended at an OD_600_ = 0.5 U in GC Broth. Then, 2-fold serial dilutions were made until at an OD_600_ = 0.0078 U in the same medium. Next, 200 µl of selected 2-fold dilutions were distributed by sextuplicate in a white-wall clear bottom 96-well assay plate (Corning). The plate was incubated in a luminescence plate reader (BioTek Synergy H1, Agilent) at 37°C for 24 hrs. The OD_600_ and luminescence data were automatically collected every 10 minutes after the cultures were agitated for 15 sec (double orbital rotation, 150 rpm). Bioluminescence output from each well was acquired for one second and presented as total relative light unit (RLU) counts/s.

### *Neisseria gonorrhoeae* axenic growth kinetics under different nutrient conditions

The bacteria were grown on GCB agar and harvested in PBSG (PBS supplemented with 7.5 mM Glucose, 0.9 mM CaCl_2_, and 0.7 mM MgCl_2_). Then, the bacteria were resuspended at an OD_600_ = 0.5 U in three different nutritional culture conditions: (i) PBSG containing 25 mM HEPES (HyClone GE Healthcare Life Sciences), (ii) RPMI medium (Genesee) with 25 mM HEPES and (iii) RPMI with 25 mM HEPES and supplemented with 10% human serum (Sigma-H4522). Then, 2-fold serial dilutions were made in each medium and 200 µl of selected 2-fold dilutions were distributed by sextuplicate in a white-wall clear bottom 96-well assay plate (Corning). The plate was incubated in the luminescence plate reader at 37°C for 24 hrs and OD_600_ and bioluminescence were acquired as described above.

### Neisseria gonorrhoeae inoculum preparation

For infection of macrophages, bacterial strains were grown from frozen stocks onto GCB agar for 16 hrs. The next day, 5 piliated colonies were pooled and streaked on GCB and were grown for another 16 hrs. On the second day, bacteria were collected and resuspended in PBSG. Bacterial numbers were determined by OD_600_ measurements. The inoculum was prepared at MOI = 5 in the media used in the infection assay. Inocula for all infections were confirmed by plating dilutions on GCB agar for CFU recovery.

### Cell culture conditions

The human monocytic cell line U937 (ATCC CRL-1593.2) was obtained from ATCC and primary human peripheral monocytic cells (PBMCs) were purchased from Sigma (HUMANPBMC-0002644). U937 cells were cultured in RPMI medium (Genesee) supplemented with 10% Fetal Bovine Serum (FBS) at 37°C and 5% CO_2_. For differentiation into mature adherent macrophages, U937 monocytes were seeded at the desired density, cultured with 10 ng/ml phorbol 12-myristate 13-acetate (PMA) for 24 hrs, followed by 48 hrs incubation with RPMI media (+10% FBS) (v/v) in the absence of PMA. For M1 and M2 polarization, U937 cells differentiated with PMA, were treated with 25 ng/ml IFNγ (BioLegend catalog #713906) and 100 ng/ml *Escherichia coli* lipopolysaccharides (LPS) for 24 hrs to induce M1 polarization or 25 ng/ml IL-4 (BioLegend catalog #766202) and 25 ng/ml IL-13 (BioLegend catalog #571102) for M2 polarization. Primary human PBMC were differentiated into macrophages (hMDMs) after culturing for 7 days with RPMI supplemented with 20% heat-inactivated human serum (Sigma-H4522).

### Quantitative analysis of bacterial replication in infection assays

Cells were seeded at 5×10^4^ cells per well in a white-wall clear bottom 96-well assay plate (Corning 3610) for infection assays. U937 monocytes were directly differentiated in 96-well plates, whereas hMDMs were differentiated first and then seeded in the plates for infection. For infection of polarized macrophages, cells cultured in RPMI (+10% FBS) (v/v) were treated with the respective cytokines (as detailed above) for 24 hrs prior to the infection. The media were replaced with either PBSG, RPMI, or RPMI supplemented with 10% human serum for the infection assays. All media were supplemented with 25 mM HEPES. For both *Neisseria* and *Legionella*, macrophages were infected at MOI = 5 and infections were carried out in six technical replicates for every condition. The 96-well plates were incubated in the luminescence plate reader at 37°C for 24 hrs (for *Ng*) or 72 hrs (for *Lp*) and bioluminescence output was obtained hourly. Light was collected for 1s per well and the data is presented as total relative light unit (RLU) counts/s.

### Microscopy analyses of infected cells

U937 monocytes seeded at 2×10^5^ cells/well were differentiated into macrophages directly on coverslips in 24-well plates (Gen Clone). Macrophages were polarized as detailed above and infected with *Ng* FA1090 strain (MOI = 5) in the indicated media for 8 hrs. To stop the infection, the coverslips were washed three times with warm PBS and fixed with 4% paraformaldehyde (PFA) (Electron Microscopy Sciences) at 4°C overnight. Subsequently, cover slips were blocked with 2% goat serum in PBS for 30 min, and an anti-*Ng* chicken IgY antibody (diluted 1:100 in PBS) was added for 1 hour at ambient temperature for detection of extracellular bacteria. Next, the coverslips were washed 3X with PBS, fixed with 4% PFA for 30min, washed 3X with PBS, permeabilized and blocked simultaneously with 0.1% TritonX-100 and 2% goat serum in PBS for 60 min at ambient temperature. After washing, an anti-*Ng* rabbit IgG antibody (Fitzgerald Ind. catalog #20-NR08) diluted 1:500 in PBS was added for 2 hours at ambient temperature to label all the bacteria in the sample. Next, the coverslips were washed 3X with PBS, and incubated at ambient temperature for 1 hour with a goat anti-chicken-Alexa Fluor488 (Invitrogen catalog #A-11039) (diluted at 1:300), a goat anti-rabbit IgG, Alexa Fluor647 (Invitrogen catalog #A21244) (diluted at 1:1000), the actin probe phalloidin-Alexa Fluor555 (Invitrogen catalog #A30106) (diluted at 1:2000), and Hoechst (Invitrogen catalog #H3570) (diluted at 1:2000). After washing, coverslips were mounted on glass slides with ProLong Gold antifade reagent (Invitrogen catalog #P36984) and were examined by fluorescence microscopy. All images were acquired with an inverted wide-field microscope (Nikon Eclipse Ti) controlled by NES Elements v4.3 imaging software (Nikon) using a 60X/1.40 oil objective (Nikon Plan Apo λ), LED illumination (Lumencor) and CoolSNAP MYO CCD camera. Image acquisition and analysis were completed with NES Elements v4.3 imaging software.

### RNA extraction and quantitative real-time PCR

The RNA extraction and quantitative real-time PCR (qPCR) were performed as previously described (18). RNA was extracted from U937 M0 macrophages or M1 and M2 polarized macrophages using the RNeasy kit from Qiagen. First-strand cDNA synthesis was performed using TaqMan Gene Expression Cells-To-Ct kit (ThermoFisher catalog: AM1728). Relative mRNA levels were determined by quantitative real-time PCR using PowerUp SYBR Green master mix (ThermoFisher). For PowerUp SYBR Green, the thermal cycling was performed under the following conditions: pre-amplification: 95°C for 10min; amplification: 40 cycles at 95°C for 10s, 60°C for 10s, and 72°C for 10s; melting: 95°C for 10s, 60°C for 60s, and 97°C for 1s. Relative quantification was performed using 2^−ΔΔCT^ method. Gene expression was normalized to *GAPDH* as reference gene. A list of primers used in this study is presented in Supplemental Table 1.

### Statistical analysis

Calculations for statistical differences were completed by Student’s T-test or multiparametric ANOVA analysis using GraphPad Prism v10 software and data are presented in the respective figures’ panels.

## Results

### Generation of bioluminescent Ng strain for bacterial growth analysis

Bioluminescence has emerged as a reliable approach for quantitative analysis of bacterial growth under various conditions, including in cellular infections (44, 45). Measurements of light output from bioluminescent bacteria allows growth kinetics analysis in a high-throughput manner with great resolution (41). Because light production requires ATP, bioluminescence also reflects the metabolic state and viability of the producers (46). We engineered a bioluminescent *Ng* FA1090 strain (*Ng* FA1090-LuxR) by inserting the LuxR operon (*luxCDABE*) from *Photorhabdus luminescens* on the *Ng* chromosome in the intergenic region of the *iga*-*trpB* locus (43) under the constitutive promoter of the gonococcal *opaB* gene using allelic exchange (Fig. 1A). The *Ng iga*-*trpB* locus has been used frequently for chromosomal complementation studies because insertion of genetic elements does not affect bacterial replication (43). The bioluminescence output by the *Ng* FA1090-LuxR strain at various densities correlated well with alternative quantitative methods – namely, colony forming unit (CFU) counts (Fig. 1B) and optical density measurement (Fig. 1C) - with a linear dynamic range of at least two orders of magnitude (Fig. 1B-C). The light output per CFU was similar (∼ 0.15 RLUs/CFU) across several population densities (Fig. 1D). In axenic liquid growth cultures with GCBL media (40), the parental FA1090 and FA1090-LuxR grew similarly indicating that bioluminescence production did not increase the metabolic burden sufficiently to affect bacterial replication (Fig. 1E). The bioluminescence output of the FA1090-LuxR strain increased exponentially during logarithmic growth phase, peaked in the early stationary phase, and subsequently decreased (Fig. 1F). Altogether, these results demonstrate that FA1090-LuxR grows with kinetics similar to the parental strain and bioluminescence production is an excellent readout in quantitative bacterial growth assays.

**Figure 1.**
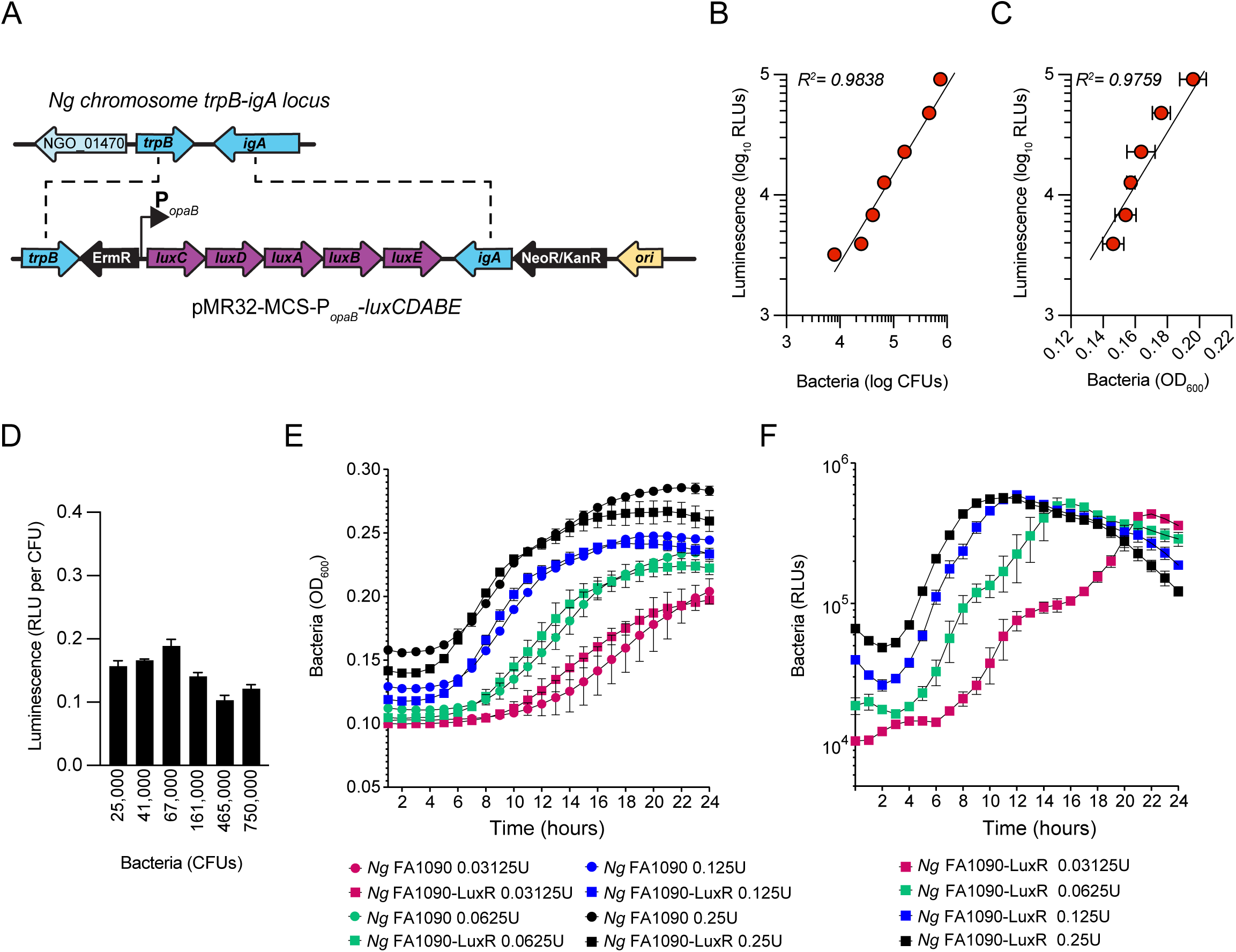
Construction and characterization of *Ng* FA1090-LuxR strain. (A) Construction of erythromycin-resistant FA1090-LuxR *Ng* strain via allelic exchange with pMR32-MCS*-luxCDABE* containing the LuxR operon cloned downstream of the constitutive *opaB* promoter (P_opaB_). (B-C) Correlation between bioluminescence output (Luminescence RLUs) and bacterial population as measured by CFUs (B) or by optical density (C) for serial dilutions prepared from bacteria grown on GCB plates overnight. (D) Bioluminescence output per CFU is stable across several bacterial population densities. (E-F) Bacterial growth in axenic cultures with GCBL medium for several serial dilutions of *Ng* FA1090-LuxR and its parental strain over the indicated time period as measured by optical density (OD_600_) (E) and luminescence output (F). (B-E) Data is presented as an average from technical triplicates ± StDev, from one of three biological replicates.

### Neisseria gonorrhoeae grows in both nutrient-rich and nutrient-depleted conditions in the presence of human macrophages

We measured the growth of the FA1090-LuxR strain in macrophages infections using bioluminescence as a readout (Figs. 2A-B). To this end, human U937 macrophages (Fig. 2A) or primary monocyte-derived macrophages (hMDMs) (Fig. 2B) were infected with FA1090-LuxR for 12 hrs in nutrients-depleted PBSG (PBS + 7.5 mM glucose) medium and bioluminescence output was measured hourly. In the absence of host cells, *Ng* does not grow in PBSG medium (17, 18) as evidenced by the progressive decline in the bioluminescence signal (Fig. 2C). Conversely, bioluminescence signal increases two orders of magnitude within 8 hrs post infection (hpi) in PBSG medium when either U937 macrophages or hMDMs were present, indicative of explosive *Ng* replication (Figs. 2A-B). Infections in nutrients replete RPMI medium slightly but statistically significantly enhanced *Ng* replication over the PBSG medium (Figs. 2A-B); moreover, RPMI medium supported limited *Ng* replication even in the absence of host cells (Fig. 2C) (17). Thus, macrophages can support robust *Ng* growth regardless of the nutrient abundance in the extracellular milieu. Next, *Ng* growth in macrophage infections was carried out in RPMI medium containing heat inactivated human serum, which further increases nutrients abundance in the extracellular environment. Addition of human serum reduced slightly the bacterial growth in U937 macrophages infections (Fig. 2A). However, addition of serum notably affected *Ng* growth in hMDM infections (Fig. 2B). Together these data indicate that the presence of host cells rather than the nutritional content of the extracellular milieu is the important factor for *Ng* replication under infection conditions. Moreover, heat-inactivated human serum restricted *Ng* growth in RPMI medium in the absence of host cells (Fig. 2C), suggestive of microbicidal activity. This serum-dependent growth restriction phenotype was more evident in hMDMs (Fig. 2B) but not in U937 macrophages infections (Fig. 2A).

**Figure 2.**
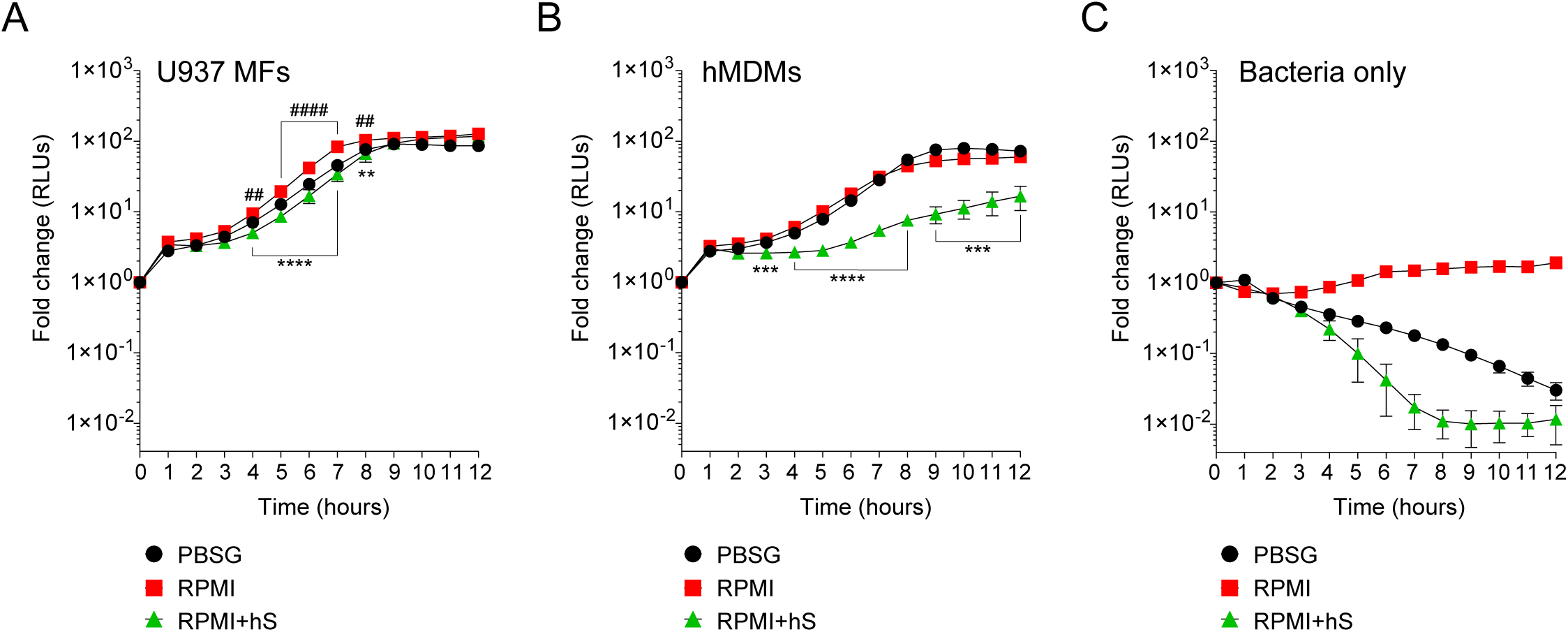
Kinetics of bacterial replication in the presence or absence of macrophages. (A-C) Quantitative analysis of FA1090-LuxR replication in the indicated media in the presence of either U937 macrophages (U937 MFs) (A) or primary human monocyte-derived macrophages (hMDMs) (B), or in the absence of host cells (Bacteria only) (C). Data is presented as fold change in bioluminescence (RLUs) from T0. Each time point is an average from six technical replicates ± StDev. Representative data from one of three biological replicates are shown. ‘hS’ – heat-inactivated human serum. Statistical differences were calculated using two-way ANOVA analysis (*, statistical significance for RPMI *vs.* RPMI+hS; ^#^, statistical significance for RPMI *vs.* PBSG. ** and ^##^, *p* < 0.01; ***, *p* < 0.001; **** and ^####^, *p* < 0.0001).

### Neisseria gonorrhoeae invasion and intracellular growth is not dependent of extracellular nutrients

Because *Ng* colonizes macrophages and can replicate intracellularly as well as on the cell surface, we investigated whether nutrients availability affects invasion or bacterial replication in either of the niches. To this end, the invasion of U937 macrophages by gonococcal colonies was measured for the first 8 hrs of infection. Statistically significant increase in macrophages invasion was observed when infections were carried out in serum-containing RPMI medium as compared to infections in PBSG (Fig. 3A) indicating that one or more factors in serum may directly or indirectly trigger the invasion process. Macrophages invasion is mediated by subversion of the actin cytoskeleton as an extensive network of cortical actin filaments forms at the plasma membrane at the invasion site and surrounds the invading colony even after internalization is complete (18). In all tested conditions, intracellular colonies were enveloped by actin filaments indicating that mechanistically the invasion process is similar under all three infection settings (Fig. 3B). However, we noted that when extracellular nutrients were present, the size of the intracellular colonies appeared larger as compared to PBSG conditions (Fig. 3B). Therefore, intracellular replication was specifically measured by imaging and 3D volumetric size analysis in assays where colonies were allowed to invade U937 macrophages for 4 hrs and then expand intracellularly for additional 4 hrs (Fig. 3C). Indeed, significant increase in the average colony size was observed in the presence of RPMI and RPMI+hS conditions. Thus, we conclude that nutrients-replete medium enhanced *Ng* intracellular replication in U937 macrophages and can enhance macrophage invasion when serum is present. Together, these data also indicate that the growth promoting effect of the nutrients in the extracellular milieu is specific for bacteria replicating intracellularly because in RPMI+hS *vs.* PBSG condition the overall bacterial growth was lower (Fig. 2A) but the intracellular growth was significantly enhanced (Fig. 3C).

**Figure 3.**
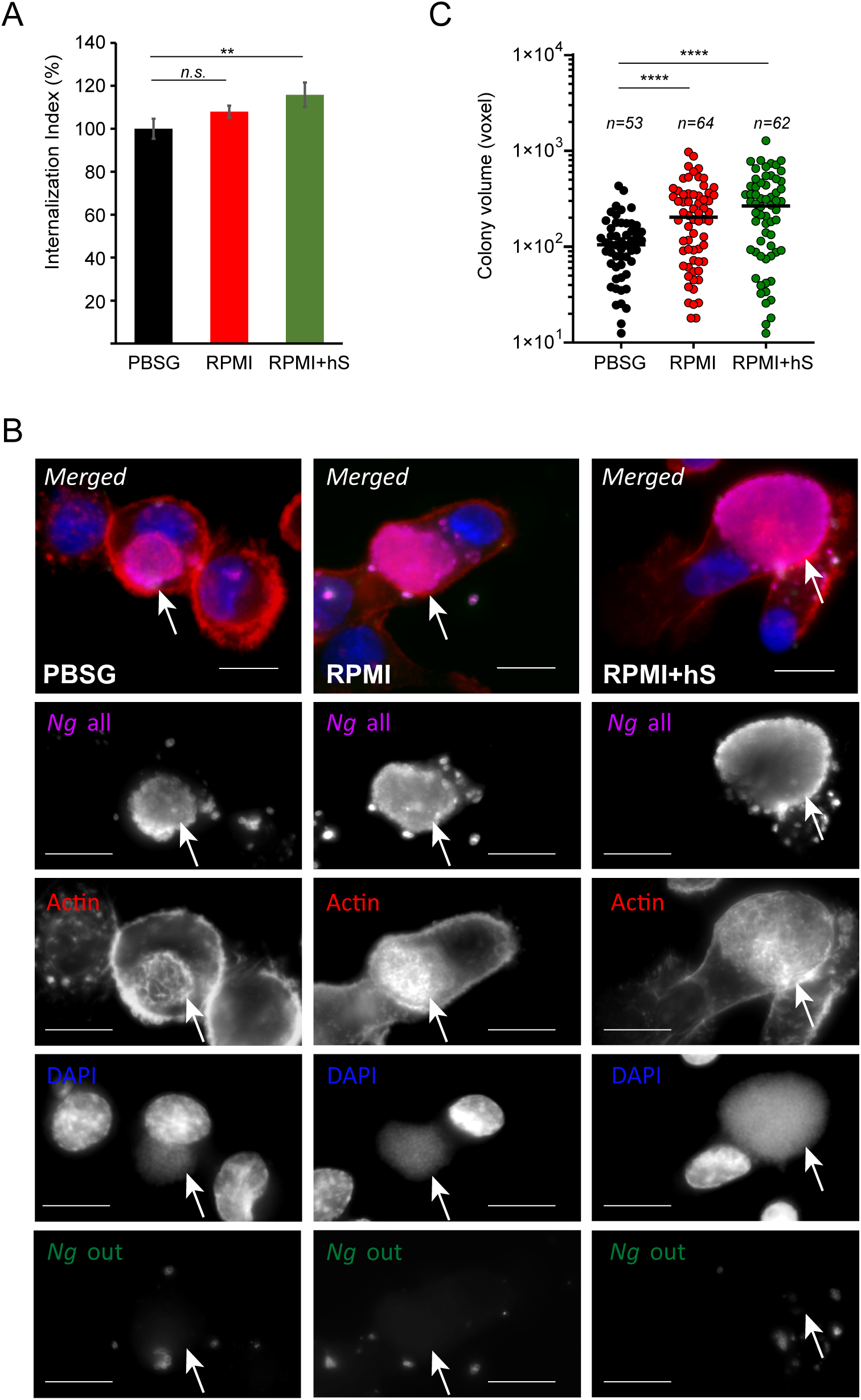
Extracellular nutrients abundance promotes gonococci invasion and intracellular growth in U937 macrophages. (A) Invasion of *Ng* colonies in macrophages cultured under nutrient-depleted (PBSG) and nutrient-replete conditions (RPMI and RPMI+hS) was determined at 8 hpi. The ‘Internalization Index’ reflects how each medium affects invasion compared to PBSG and was calculated for each condition by dividing the percentage of internalized microcolonies from the ‘RPMI’ and ‘RPMI+hS’ conditions by the percentage of internalized microcolonies from the ‘PBSG’ condition, which was then multiplied by 100. Graphs show averages of three biological replicates ± StDev. (B) Representative micrographs showing intracellular microcolonies at 8 hpi as determined by inside/out staining, where infections were carried out under nutrients-deplete (PBSG) or nutrients-replete (RPMI or RPMI+hS) conditions. Individual channels of the merged images are shown in grayscale. Arrows denote individual intracellular colonies. Bar = 10 μm. (C) Size measurements of individual intracellular colonies under the indicated growth conditions using 3D object volume microscopy measurements (in voxels). Each data point is derived from a single colony. The number of colonies included in the analysis is indicated for each condition. Means for each object population is shown. Representative data from one of three biological replicates are shown. (A, C) Statistical significance was determined via unpaired T-test analysis (*n.s.*, not significant; **, *p* < 0.01; ****, *p* < 0.0001).

### Macrophages in different immune activation states support Neisseria gonorrhoeae replication and invasion

To assess whether the phenotypic state of the macrophage impacts *Ng* capacity to colonize, invade or replicate with it, U937 macrophages were polarized in pro-inflammatory M1-like state by stimulation with IFNγ and *E.coli* LPS, or in the tolerogenic M2-like state by treatment with IL-4 and IL-13 or were left in the M0 state (47). Successful polarization in the various phenotypic states was confirmed by the upregulation of state-specific cellular markers – *IRF1* and *IL6* for M1 and *IL10* for M2 (Fig. 4A). Microbiocidal functions are enhanced in M1 macrophages (48), therefore we investigated whether M1-polarized U937 macrophages under the conditions we used restrict the replication of the vacuolar pathogen *Legionella pneumophila* as it would be expected (49, 50). Indeed, M1-polarized U937 macrophages significantly restricted *Legionella* growth as compared to M0 controls (Fig. 4B). However, M1-polarized U937 macrophages failed to restrict *Ng* growth irrespective of the cell culture media (PBSG, RPMI or RPMI+hS) used in the infection assays (Figs. 4B-C). Because IFNγ-dependent cell autonomous defenses mainly target intracellular pathogens (50, 51, 52, 53, 54), the lack of growth restriction by M1 macrophages might indicate that either intracellular gonococci can interfere with microbiocidal activities of M1 macrophages or perhaps enhanced replication by surface-associated bacteria compensates for a potential decrease in the intracellular population. To distinguish between those possibilities, we directly measured *Ng* capacity to invade and grow within M1 macrophages (Figs. 5A-C). At 8 hpi, the number of gonococcal colonies inside M1 macrophages was reduced by ∼40% for U937 cells (Fig. 5A) and ∼60% for hMDMs (Fig. 5B), indicating that IFNγ-priming reduces invasion, or a fraction of intracellular colonies are eliminated. When we measured expansion of gonococci intracellular colonies housed by the different macrophage types, the data revealed similar growth rates indicating that M1-polarized macrophages likely interfere with gonococcal invasion and not with intracellular replication (Fig. 5C). Because the host actin nucleating factor FMNL3 mediates gonococcal invasion in macrophages (18), we compared the amount of *FMNL3* transcripts in polarized macrophages using qPCR analysis (Fig. 5D). *FMNL3* expression in M0 and M2 U937 cells was indistinguishable, however in M1 U937 cells the mRNA levels were reduced by 60% (Fig. 5D). Thus, M1-porization results in decrease of a host factor mediating gonococcal invasion, which potentially might explain the defect in M1 cells. If lowered FMNL3 expression in M1 reduced gonococcal invasion, it would be expected that overexpression of FMNL3 from a heterologous promoter should restore invasion in M1 macrophages. To this end, we used a pool of U937 macrophages in which ∼50% of the cells stably overexpress V5-tagged FMNL3 from a CMV promoter to measure gonococcal invasion (18) (Fig. 5E). Invasion was equally reduced under M1-polarizing conditions regardless of whether macrophages overexpressed FMNL3-V5 or not (Fig. 5F), indicating that lower FMNL3 expression alone does not account for the invasion blockade. The FMNL3-V5 allele is functional and has previously complemented the invasion defect observed in FMNL3 KO macrophages (18). Thus, we conclude that: (i) M1-polarization reduces *FMNL3* expression and blocks gonococcal invasion by a mechanism that cannot be overcome by restoration of FMNL3 abundance; (ii) macrophages support robust intracellular as well as extracellular gonococcal replication regardless of the phenotypic polarization state and the availability of extracellular nutrients; (iii) increased replication of surface-associated bacteria likely compensates for the decreased intracellular bacterial population brought by the invasion blockade under M1-polarization conditions. Taken together, our data demonstrate that gonococci have evolved mechanisms to colonize successfully and replication in association with macrophages regardless of their polarization state.

**Figure 4.**
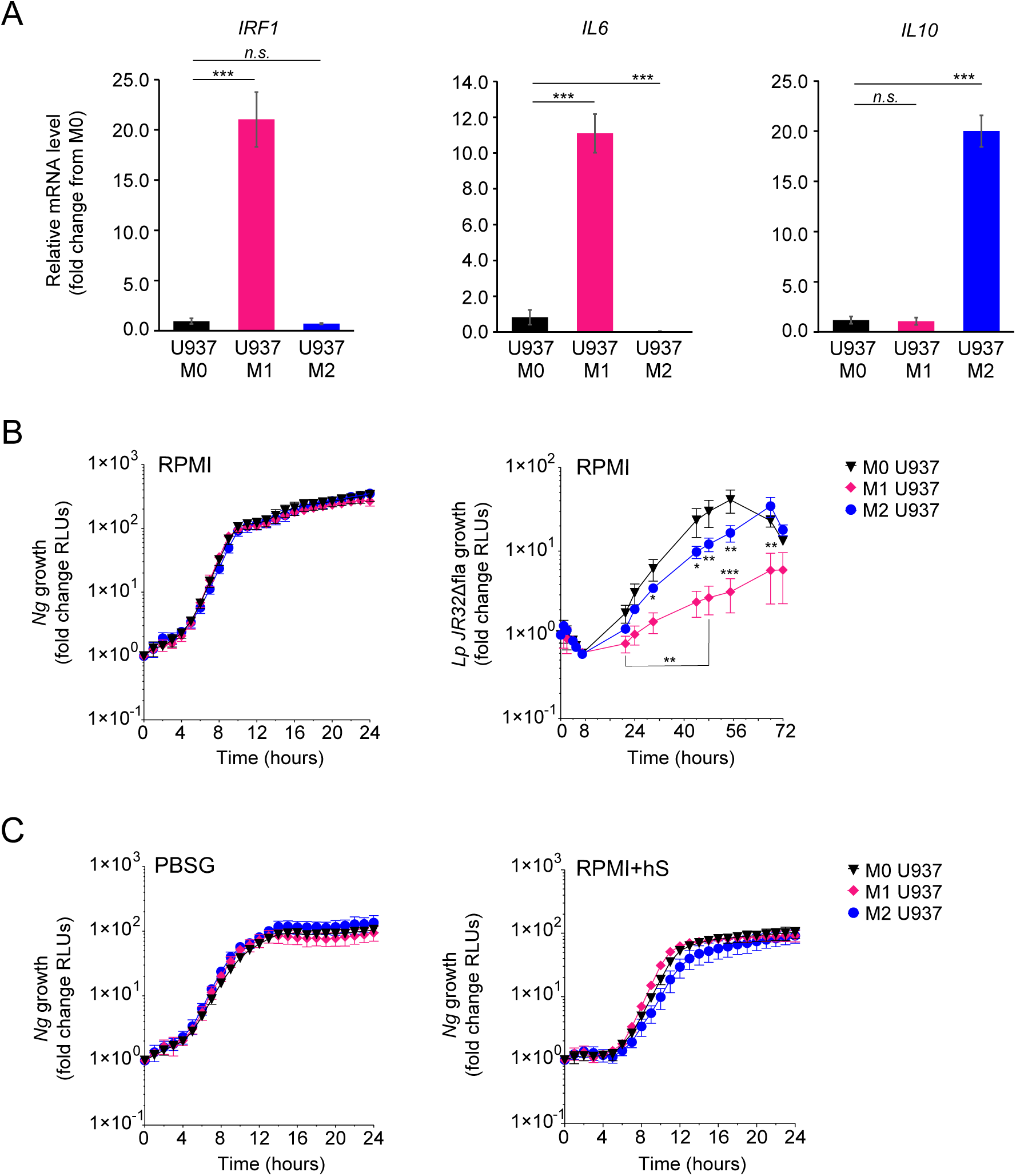
Kinetics of *Ng* replication with polarized U937 macrophages. (A) U937 macrophages polarization in M0, M1 and M2-like states was validated by qPCR analysis for expression of different markers associated with each phenotypic state – *IRF1* and *IL6* for M1 and *IL10* for M2. Graphs represent gene mRNA expression relative to M0 (fold change). (B-C) Kinetics of bacterial replication supported by M0, M1 and M2 macrophages under conditions of nutrient abundance or restriction. Growth is presented as fold change in bioluminescence for each condition from T_0_. Data from *Ng* and *Lp* infections in RPMI are shown in (B) and data from *Ng* infections in PBSG or RPMI+hS are shown in (C). (B-C) Each data point on the graphs represents an average from technical triplicates ± StDev. Representative data from one of three biological replicates are shown. (A) Statistical significance was determined via unpaired T-test analysis (*n.s.*, not significant; *\**\*\*, *p* < 0.001). (B) Statistical differences were calculated using two-way ANOVA analysis (*, *p* < 0.05; **, *p* < 0.01; ***, *p* < 0.001).

**Figure 5.**
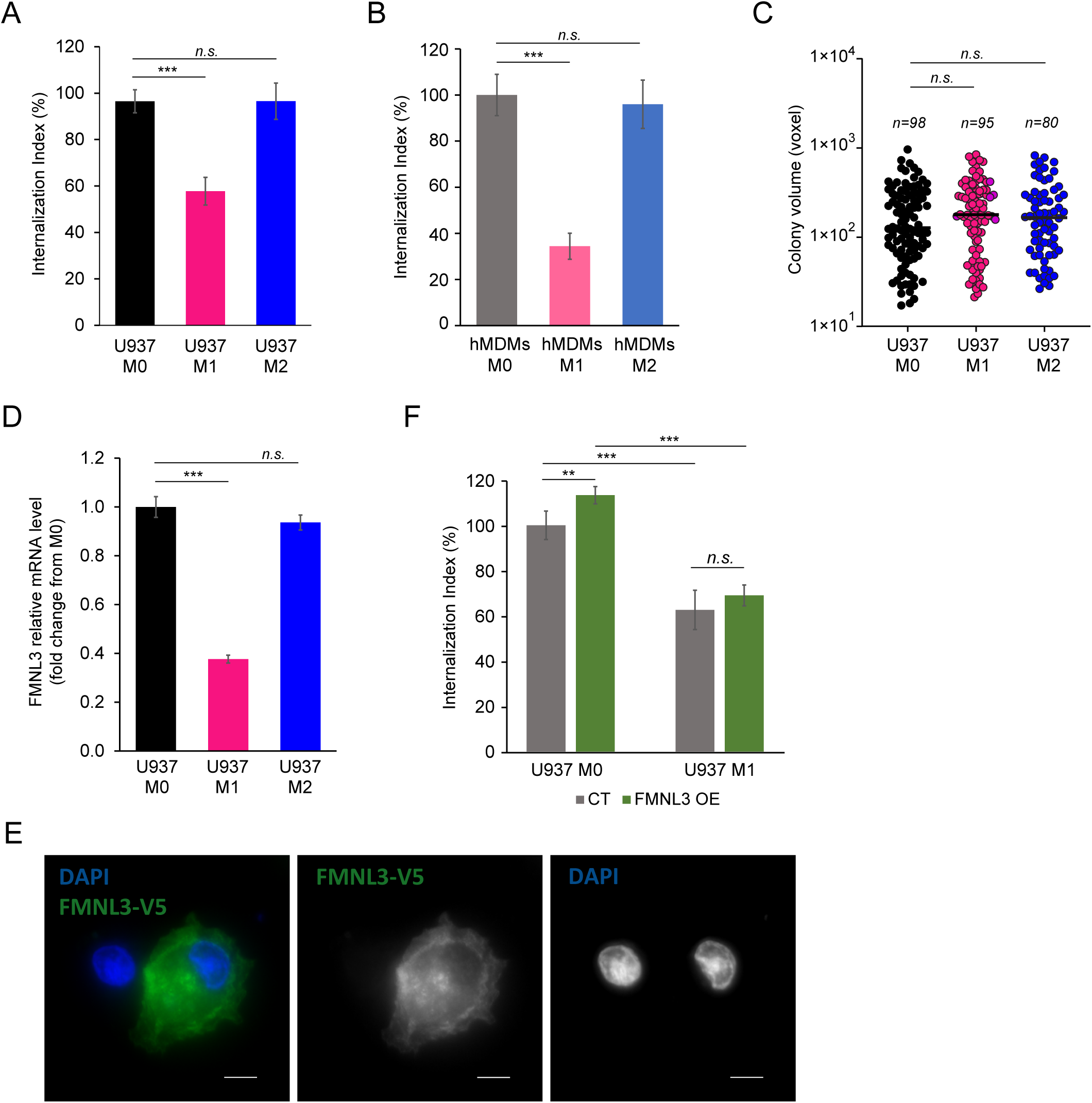
*Ng* invasion and intracellular replication in polarized macrophages. (A-B) Invasion of *Ng* colonies in polarized macrophages was determined at 8 hpi. The ‘Internalization Index’ reflects how each phenotypic state affects invasion compared to M0 and was calculated for each condition by dividing the percentage of internalized microcolonies from M1 and M2 conditions by the percentage of internalized microcolonies by M0 macrophages, which was then multiplied by 100. Data from infections of polarized U937 macrophages (A) and hMDMs (B) are shown. Graphed data represent averages of three biological replicates ± StDev. (C) The sizes of intracellular colonies harbored by polarized U937 macrophages as determined by 3D object volume microscopy measurements (in voxels). Each data point is derived from a single colony. The number of colonies included in the analysis is indicated for each condition. Representative data from one of three biological replicates are shown. (D) Gene expression analysis for *FMNL3* in polarized U937 macrophages as determined by qPCR. Graphed data represent *FMNL3* expression under different conditions relative to expression in M0 (fold change). (E) Representative micrograph showing FMNL3-V5 expression in a pool of U937 macrophages that either overexpress the allele (right cell) or not (left cell). Individual channels of the merged images are shown in grayscale. Bar = 5 μm. (F) Gonococcal invasion of a heterogenous pool of M0 or M1 polarized U937 macrophages that either express FMNL3-V5 (OE) or not (CT). The ‘Internalization Index’ was calculated at 8 hpi and is normalized to the percentage of colonies internalized by M0 CT cells, which was set at 100%. Graphs show averages of three biological replicates ± StDev. (A-D, F) Statistical analyses were completed by unpaired Student’s T-test (*n.s.*, not significant; **, *p* < 0.01; ***, *p* < 0.001).

## Discussion

Lack of a protective immunity against reinfection represents one notable aspect of gonococcal infections that hampers vaccine development. The weakened immunological response is a result of extensive host adaptation and development of multiple innate and adaptive immunosuppressive mechanisms (55, 56, 57). The subversion of mucosal macrophages – the tissue sentinels in the genitourinary tract (8, 19) – into putative cellular niches and immunoregulators in gonococcal infections is noted as an important knowledge gap (56, 57). Once gonococci cross the urogenital epithelium and invade the submucosal space, the bacteria gain access to tissue-resident as well as monocytes-derived macrophages recruited to the infection site. Previous studies showed that gonococci can promote an M2-like polarization state in human macrophages which is incapable of inducing CD4+ T cell proliferation (58). Gonococci elicit the production of pro- as well as anti-inflammatory cytokines when cultured with macrophages *in vitro*. However, IL10 secretion was induced at much lower bacterial loads as compared to IL6 (58), indicating potential skewing towards a more anti-inflammatory response, which is consistent with the prevalent asymptomatic nature of the infection. Here we investigated whether and to what extent macrophage polarization affects the cell capacity to function as a niche for bacterial replication. When gonococci colonize macrophages, bacterial replication takes place either in a surface-associated or within an internalized microcolony (18). Despite potent microbiocidal capacity towards intracellular bacteria, our data demonstrates that M1 macrophages could not restrict *Ng* growth but did reduce invasion. These observations indicate that *Ng* replication likely takes place preferentially in surface-associated colonies because of reduction in the number of intracellular microcolonies. These data further support our original premise that dual niche occupancy enhances evasion of cell autonomous host defenses (18). Moreover, gonococci intracellular replication in M1 and M0 macrophages was indistinguishable despite a robust restriction of *Legionella* intracellular replication after M1 polarization. This phenotype provides strong evidence for the existence of *Ng*-encoded mechanisms directed towards evasion of microbiocidal activities of M1 macrophages. The unique membrane-bound organelle harboring the intracellular gonococcal colony does not fuse with lysosomes and lacks endosomal markers (18), and our data demonstrates that even under M1 polarization conditions the vacuole protects the bacteria from host microbiocidal activities. Disruption of pathogen-occupied vacuoles by IFNγ-inducible gene products can trigger cytosolic cell autonomous defense responses that culminate in either host cell death or autophagolysosomal clearance (54, 59, 60). However, we did not observe signs of either in M1 macrophages invaded by *Ng* (data not shown) indicating that the vacuolar compartment is intact and supported bacterial replication. However, gonococci have been shown to interfere with apoptotic responses, thus a blockade of IFNγ-dependent host cell death response may also play a role for intracellular survival in M1 macrophages. The immune evasion mechanisms facilitating intracellular replication in M1 macrophages warrants further investigation. It is notable that under all polarization conditions tested, gonococci replication rates intracellularly and overall (intracellularly + extracellularly) were comparable, indicating that *Ng* is well adapted to exploit macrophages for growth regardless of their polarization state. This was not the case for *Legionella* – an accidental human pathogen – which could not overcome the antimicrobial defenses of M1 macrophages.

Our data demonstrated that M1 polarization reduced invasion but not intracellular replication. Notably, stimulation with IFNγ/LPS decreased expression of FMNL3 – an actin nucleator from the formin family that mediates actin polymerization necessary for gonococcal invasion (18). Overexpression of FMNL3 under heterologous promoter did not increase invasion in M1 macrophages; therefore, other regulators upstream or downstream of FMNL3 are also affected as a result of M1 polarization. Indeed, we have shown that multiple formins or myosin motors regulate gonococcal invasion (18).

The present study also investigated the impact of extracellular nutrients on gonococcal replication with macrophages. In the absence of macrophages, limited gonococcal growth is observed in nutrient-replete medium (RPMI), whereas nutrient-depleted (PBSG) medium did not support gonococcal replication (Fig. 2B) (17, 18). When either primary or immortalized macrophages were present, gonococci population overall increased 2 orders of magnitude within 8 hpi regardless of the culture media used in the assays. Notably, when heat-inactivated human serum was provided, gonococcal growth with hMDMs was delayed and reduced as compared to serum-free infections, indicating that a serum-associated activity either directly or indirectly might interfere with gonococcal growth. Direct interference with bacterial replication by human serum was also observed in the absence of macrophages. Moreover, intracellular gonococci were protected from the anti-microbial activity of the human serum in macrophage infections and intracellular growth increased under those conditions. Gonococci invade U937 macrophages faster as compared to hMDMs (18), thus the number of intracellular bacteria as percentage of the total population is also higher, which explains why the bactericidal effect of human serum, which targets surface-associated and extracellular bacteria, is pronounced in hMDM infections. The increase in intracellular growth likely compensates partially for the decrease in extracellular replication when serum is present, and we also observed increased invasion when human serum was present in the culture media. Collectively, our data demonstrates intracellular gonococci replication is primarily affected by nutrient content of the extracellular milieu, where intracellular growth occurred under all conditions but was significantly enhanced when nutrients were present.

