## Supplemental Table 1 for "Characterization of *Neisseria gonorrhoeae* colonization of macrophages under distinct polarization states and nutrients environment"

| **Gene name** | **Forward primer** | **Reverse primer** |
|---|---|---|
| *IRF1* | CTCTCACCAAGAACCAGAGAAA | GAAGGTATCAGGGCTGGAATC |
| *IL6* | GATGAGTACAAAAGTCCTGATCCA | CTGCAGCCACTGGTTCTGT |
| *IL10* | TGCCTTCAGCAGAGTGAAGA | GCAACCCAGGTAACCCTTAAA |
| *FMNL3* | TGGCATATACCACCCATCTCT | GTTCAAGGGTCCCCTACTCC |
| *GAPDH* | AGCCACATCGCTCAGACAC | GCCCAATACGACCAAATCC |
